# Velocities of Hippocampal Traveling Waves Proportional to Their Coherence Frequency

**DOI:** 10.1101/2024.07.21.604460

**Authors:** Gadi Goelman, Tal Benoliel, Zvi Israel, Sami Heymann, Juan Leon, Dana Ekstein

## Abstract

Cortical traveling waves, characterized by their spatial, temporal, and frequency attributes, offer insights into active regions, timing, frequency, and the direction of activity propagation. Recent evidence suggests that these waves’ directionality and spatiotemporal extent encode cognitive processes, yet the encoding mechanism related to their frequency remains unclear.

We explore the hypothesis that coherence frequency dictates the velocity of wave propagation. Using nonlinear coherence analysis to compute propagation pathways among four local-field-potential signals collected along the human hippocampus’s long axis, we present evidence that the coherence frequency of traveling waves encodes temporal communication aspects. Unlike linear analyses, which may overestimate velocities due to bidirectional flow when considering multiple pair coherences, nonlinear analysis treats pathways as holistic units with affectively unidirectional flow, making it more suitable for calculating wave velocities. Our findings reveal that propagation velocities along the hippocampus at low frequencies (<∼35Hz) demonstrate a linear dependency on frequency, with an increased slope at higher frequencies, suggesting different underlying mechanisms. Although observed within the hippocampus, these findings capture a dependency between frequency and velocity of traveling waves which may be applicable to other cortical areas as well.

## 1. Introduction

Traveling waves that are based on coherence [1, 2], have been observed in multiple regions in humans[3, 4] and animals [5, 6], at both small and large scales. They have been suggested to elucidate the coordination of neural activity between regions, the mechanism of large-scale neural organization [1, 2], and the brain’s ability to dynamically adapt to behavioral and cognitive changes[7, 8]. Recently, it has been demonstrated that in episodic memory, measured directly in humans, the directions of propagations encode different memory processes, suggesting a link between directionality and the timing of regional activity with cognition and behavior [9]. Since traveling waves frequency, spatial and temporal aspects are thought to indicate which region is active, at what time, at what frequency, and in which direction activity is propagating, it is crucial to understand what determines the velocity of propagation.

Inspection of time-frequency wavelet plots of coherence between local-field-potential time-series signals consistently revealed that the width of coherence windows, assumed to represent periods during which communication takes place[10, 11], exhibits broader characteristics for lower frequencies and narrower features for higher frequencies (for example, see Figure 1 in[12]). This consistent observation strongly implies a proportional relationship between the size of time windows and the associated wavelength.

**Figure 1:**
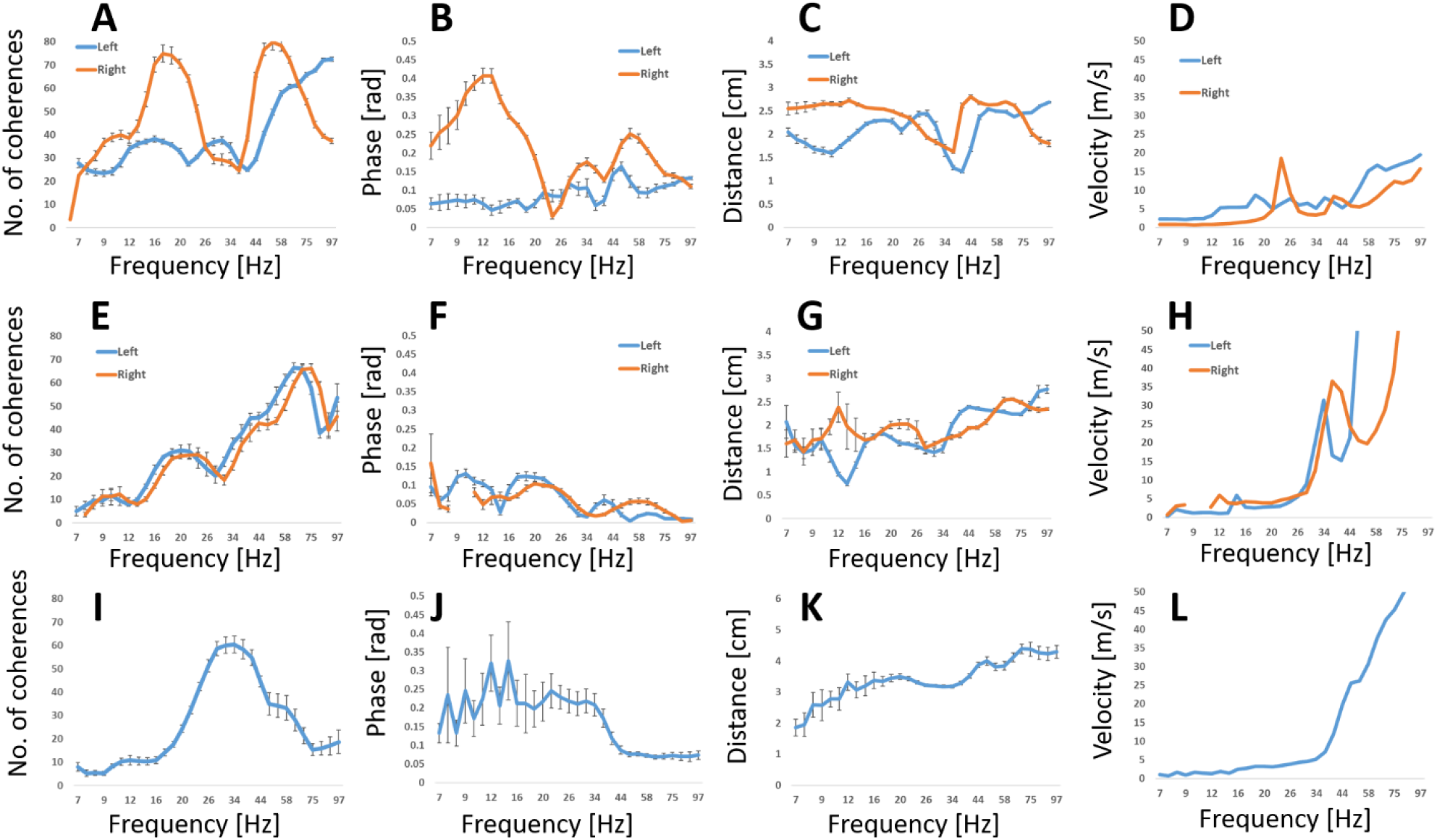
Pair coherence parameters as a function of frequency. **Top row** – Results for subject 1 left and right hippocampi. **Middle row** – Results for the left and right hippocampi of subject 2. **Bottom row** – right hippocampus of subject 3. **Left column (A, E and I)**: Normalized number of coherences. Normalization was to the maximum available combination number (105). **Second left column (B, F, J)**: Coherence phase in radians, **Second right column (C, G, K)** : Coherence distance in cm, and **Right (D, H, L):** velocity of transmission in m/s. The mean values over sets ± standard errors are shown.

This suggests that communication facilitated by coherence is governed by the duration of these time-frequency windows, thus exhibiting frequency-dependent characteristics. Consequently, based on findings reported by others [13, 14], we hypothesize that coherence frequency dictates the velocity of traveling wave propagation, so that higher frequencies correspond to faster propagations, whereas lower frequencies correspond to slower propagations.

To this end, we conducted an analysis of traveling waves within local field potential (LFP) signals extracted from the human hippocampus in neurosurgical epileptic subjects undergoing stereo-electroencephalography (SEEG) electrode implantation. In contrast to previous studies employing various linear methods for measuring traveling waves, we utilized a novel multivariate nonlinear analysis method previously applied to functional MRI signals [12, 15-20]. This nonlinear analysis leverages mutual coherence (simultaneous coherence) among four time-series (i.e., LFP signals, each from a distinct location), along with the temporal delays at each frequency between these signals, to establish their temporal order and infer directed pathways. This approach is particularly suitable for studying traveling waves as it treats pathways as holistic units rather than mere combinations of their constituent parts and assumes effectively unidirectional flow along these pathways.

Utilizing coherence to estimate transmission times and velocities relies on the assumption that the time of transmission is proportional to coherence phase over frequency, representing the time-lag between regions. However, the bidirectional nature of many cortical and hippocampal connections can lead to coherence with zero or close to zero phase lag (zero-lag synchrony) [21]. Consequently, employing linear analyses by considering multiple pair coherences, each including a mixture of connections, to track wave propagation, may lead to underestimated transmission times and, consequently, overestimated velocities. This discrepancy could manifest in varying velocities for regions where the linear assumption is approximately valid versus those where it is not, potentially contributing to the inconsistency in reported velocity values (see discussion). In contrast, the underlying assumption in the calculation of pathways by multivariate analysis is unidirectional flow between its regions. Consequently, wave propagation velocity is calculated only for pathways of effectively unidirectional flow. This renders multivariate analysis most suitable for accurately measuring transmission times from coherence phases. Results from both linear and nonlinear analyses are compared to elucidate and illustrate this issue.

Cortical waves along the hippocampus can reasonably be approximated as a one-dimensional pathway structure, enabling the estimation of propagation distances necessary for determining transmission velocities. Thus, we focused on the rare cases involving subjects with SEEG electrodes positioned along the longitudinal axes of the hippocampus. We included data from five hippocampi of three patients undergoing SEEG in our analysis. To avoid dependence on specific cognitive processes that may occur at particular frequencies, we utilized large datasets measured during spontaneous activity (resting-state data), which is assumed to encompass various cognitive and non-cognitive processes.

## 2. Results

Three subjects with drug-resistant epilepsy undergoing intracranial EEG monitoring, which included depth electrodes inserted along the hippocampal longitudinal axis as part of their pre-surgical evaluation, were included in the study. For two subjects, the SEEG electrodes spanned the anteroposterior axis of both hippocampi, and for one subject, the SEEG electrode spanned the anteroposterior axis of only the right hippocampus. Consequently, the analysis was performed on data recorded from five hippocampi over approximately three days.

For bivariate (pair coherence) calculations, we used the phase-lag-index (PLI) [20, 21], and for multivariate coherence four-node pathways, we used the pathway-index (PWI) [10-16, 22] (see method). Both calculations employed wavelet analysis with custom-developed software. To ensure accurate and reliable results while minimizing the influence of abnormal brain activity, we carefully selected spontaneous activity data occurring at least 1 minute after and 1 minute before seizures, as determined through meticulous visual inspection of the electrodes’ signals and video recordings by an expert epileptologist. All the data used in the analysis were recorded during the arousal state, identified using scalp EEG and video recordings, with subjects lying in their beds and not engaging in any specific activity.

To derive meaningful results, the PLI and PWI were computed by summing over 500 1-second time windows. Between eight and fourteen sets were selected at different times during recording, each consisting of 500 time windows, independently calculated, and the results averaged. To avoid bias introduced by the selection of specific time-series (electrodes’ contacts), we considered all possible combinations of pairs and four-node pathways, comprising 105 pair combinations and 1365 pathway combinations (since each pathway involved four contact signals). This approach ensured an equal chance of occurrence for all contacts in the PLI and PWI calculations.

To assess the potential impact of spikes, simulations were conducted with spiked data, and the aforementioned calculations were repeated. Our results demonstrated that the spikes fell outside the wavelet confidence region (defined in Torrence and Compo [18, 19]), which were not included in the calculations. This confirmed that the findings were not influenced by wavelet-filtering artifacts [19, 20].

Figure 1 shows the pair coherences across frequency for all five hippocampi. The velocity was determined by the formula: Euclidean distances ⁄time, with time defined as phase⁄frequency. Notably, the phase values, representing the time lag between time-series, generally remained low, often below

0.2 radians. This led to shorter times and consequently higher velocities particularly at high frequencies. As highlighted in the introduction, the expected bidirectional nature of communications resulted in an underestimation of the phase values, contributing to elevated velocity values. Examining individual subjects: subject 1, with relatively higher phase values at high frequencies than other hippocampi, showed a more gradual increase in velocity at high frequencies. Both hippocampi of subject 2 and the hippocampus of subject 3 exhibited a shift in velocity increase in high frequencies, attributed to the low phase values at these frequencies.

Figure 2 shows the four-node coherences of the four-node pathways across frequencies for all five hippocampi. These pathways depict propagating waves along the long axis of the hippocampus, representing a cohesive unit of interconnected hippocampal regions. Like Figure 1, it presents the mean values ± standard errors, computed over sets, for various parameters. In contrast to Figure 1, the phase values were higher, resulting in significantly lower velocities (note the ten times smaller y-axes).

**Figure 2:**
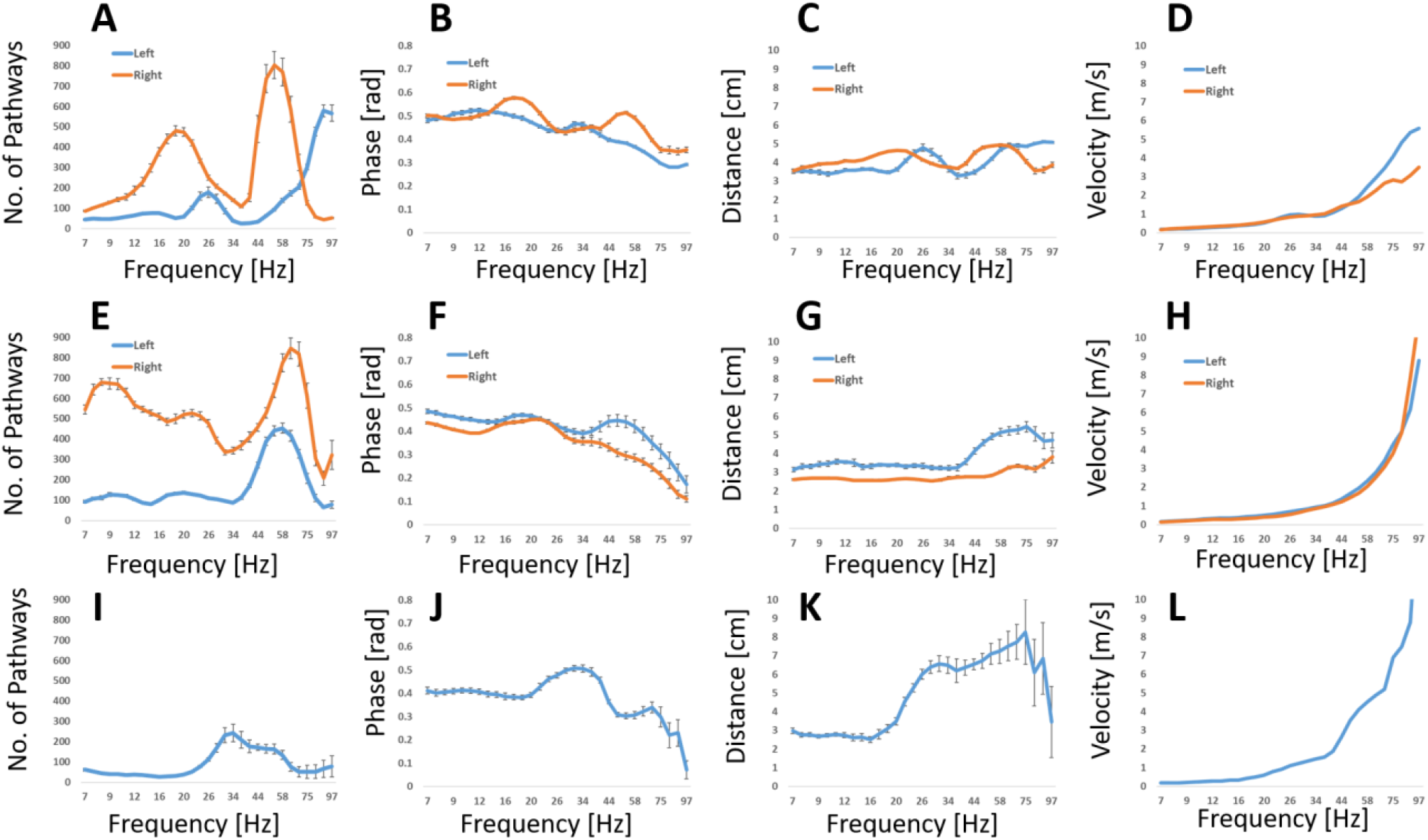
Mutual coherences (propagating waves) parameters as a function of frequency. **Top row** – Results for subject 1 left and right hippocampi. **Middle row** – Results for the left and right hippocampi of subject 2. **Bottom row** – right hippocampus of subject 3. **Left column (A, E and I)**: Normalized number of coherences. Normalization was to the maximum available combination number (1365). **Second left column (B, F, J)**: Coherence phase (mean of the three phases along pathways) in radians. **Second right column (C, G, K)**: Coherence distance (mean of the three distances along pathways) in cm, and **Right (D, H, L):** velocity of transmission in m/s. The mean values over sets ± standard errors are shown.

Generally, phase values decrease with frequency, while transmission distances and velocities increase with frequency. The velocities of all hippocampi showed a moderate increase with frequency for alpha to beta frequencies and a sharp increase at gamma frequencies. For alpha to beta frequencies, velocity followed an approximate linear dependency of: C(*v*) ∼ °.°2. *v* with a constant in units of meters, which gives spatial frequency of 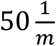 and spatial wavelength of 2cm which approximate pathway’s length at those frequencies. The frequency where the linear dependency changed to a sharp increase varies between hippocampi: approximately 35Hz for both hippocampi of subject 1, 20Hz for the right and 44Hz for the left hippocampi of subject 2, and around 20Hz for the hippocampi of subject 3. These values suggesting a moderate linear dependency of velocity on frequency for alpha to beta frequency bands, but a sharper dependency for the gamma band.

## 3. Discussion

This study aims to delve into the information encapsulated within the frequency of neuronal oscillations of traveling waves and to test the hypothesis that travelling wave velocity is proportional to the wavelength of oscillation. Due to bidirectional communication in most hippocampal and cortical pair-connectivity, employing multiple pair coherences (by using bivariate analysis) to estimate wave propagation velocity might introduce bias and lead to overestimation. This is because the pair coherence between LFP signals could encompass a blend of both unidirectional and bidirectional connections. We surmise that the reported values for hippocampal wave propagation velocities at theta frequency [13], akin to those we observed for pair coherence (Figure 1), may be inflated due to the inclusion of bidirectional connections. To mitigate this limitation, we devised a novel multivariate analysis approach that assesses mutual coherences among four coupled ensembles, constituting a pathway through which the wave propagates. This non-linear coherence characterizes the interaction between these ensembles and is tailored for unidirectional transmission along the pathways. By incorporating four-node pathways, our analysis ensures calculations specifically account for unidirectional wave propagation rather than a mixture of different connections.

Analyses were performed on SEEG electrodes signals collected from the hippocampi of neurosurgical epileptic subjects. Specifically, the data were derived from depth electrodes strategically placed along the long axes of the hippocampus, providing an optimal dataset for testing the proposed hypotheses. The choice of the hippocampus was motivated by its anticipated one-dimensional flow, enabling the estimation of transmission distances and consequently, the calculation of transmission velocities.

Electrode placement allowed for the collection of data from five hippocampi in three subjects, each recorded for approximately three days. This extended recording period enabled averaging over multiple time-windows, minimizing the impact of uncontrolled fluctuations on the analysis. The choice of spontaneous activity (resting-state) data over stimulated data, with the latter tuned to frequencies operated by the specific stimulation used, guarantees a broad band of frequencies that are expected during spontaneous activity. The fact that similar findings were observed in all five hippocampi, regardless their different health state, implies that the linkage between frequency and velocities reflects a physiologic, rather than pathologic, attribute of hippocampal information processing.

Our primary findings reveal that the velocity of transmission, calculated by pair and by four-node coherences, increases with frequency for all hippocampi. However, and as anticipated, the values are much higher for pair coherence and likely are biased by bidirectional communications. In contrast, velocity values obtained by the multivariate analysis are in line with previous publications and demonstrate different dependency for low and high frequencies. This proposition aligns with the prevailing understanding that there is no singular mechanism dictating communication speed in the brain. Instead, communication is influenced by a combination of factors, including the type of connection, myelination, synaptic transmission, and neural oscillations. Therefore, while our proposal suggests a general principle of communication through neural oscillations, various mechanisms may be in play influencing the different results obtained for lower (<35Hz) and higher (>35Hz) frequencies.

Regarding the speed of wave propagation, using EEG simulation, stable wave propagation velocity was observed within a biologically plausible range akin to axonal conduction speeds of 1–10 m/s [26].

Furthermore, in the prefrontal cortex of monkeys, the velocity of traveling waves within theta, alpha, and beta frequency ranges exhibited a linear increase with frequency, ranging from approximately 0.2 m/s to 0.5 m/s [14], consistent with the trends observed in our findings illustrated in Figure 2.

Additionally, in a separate study focusing on memory processing in humans at approximately 9 Hz, cortical propagating waves were found to travel at approximately 1 m/s[9]. These varied velocities may arise from the utilization of linear analyses, which might be suitable only in specific systems[12] or under specific stimuli. Additionally, estimating cortical propagation distances is challenging since the exact propagation pathways are complex and difficult to identify with limited coverage by SEEG electrodes.

Therefore, we propose that employing multivariate nonlinear analysis on data collected along the hippocampus is likely to yield more accurate results.

Our observation of a sharp increase in velocity for high frequencies (>35 Hz) suggests that low and high frequencies exhibit distinct dependencies on frequency. This observation aligns with the prevailing notion that neuronal oscillations in low versus high frequencies involve different underlying mechanisms. While oscillations at low (<30 Hz) frequencies are deemed crucial in neuronal communication and processing [22, 23], high-frequency oscillations are primarily attributed to local activity, often stemming from broadband multi-unit spiking activity [24, 25]. Consistent with these disparities, studies have shown that memory encoding and retrieval processes in whole-brain human connectivity tend to desynchronize for fast gamma (30–100 Hz) and synchronize for slow theta (3–8 Hz) during encoding and retrieval [26]. Moreover, it has been demonstrated that trial-by-trial fMRI BOLD fluctuations positively correlate with trial-by-trial fluctuations in high-EEG gamma power (60–80 Hz) and negatively correlate with alpha and beta power, suggesting distinct mechanisms for different oscillation frequencies [27].

Our results may provide insight into the temporal dynamics governing brain circuits, and raises the question of how varying velocities of information transfer within a network may influence the network’s computational qualities. For instance, using magnetoencephalography human data at resting state, a loop composing of higher frequency bands (alpha and betta) posterior-to-anterior information flow dominated by regions in the visual cortex and posterior default mode network was observed, and an anterior-to-posterior flow in the theta band, involving mainly regions in the frontal lobe that were sending information to a more distributed network [28]. Similarly, studies in monkeys have demonstrated that theta and gamma oscillations in the visual system promote information flow in the feedforward direction during bottom-up processing, while beta oscillations facilitate feedback interaction during top-down processing[29]. Building upon our findings, we interpret these observations as comprehensive feedback cycles, encompassing a faster bottom-up flow facilitating processing in higher regions, alongside a slower top-down flow, enabling balanced bottom-up and top-down circuits.

In summary, utilizing nonlinear coherence analysis, which inherently assumes unidirectional flow along its pathways, applied to data collected along the long axes of the hippocampus with its expected one-dimensional flow, we present evidence indicating that the coherence frequency of traveling waves along the hippocampus encodes the temporal aspects of communication. Specifically, velocities at low frequencies (<∼35Hz) exhibit a linear dependency on frequency with a spatial frequency of approximately 50 1/m, while at higher frequencies, the velocities show stronger dependence on frequency. These sharp transitions in velocities suggest different underlying mechanisms. Although these findings were observed within the hippocampus, we propose that they are applicable to other cortical areas as well.

### Limitations

Our study utilized data collected from unhealthy hippocampi, each affected by varying degrees of disease duration and severity. While efforts were made to select data exhibiting no abnormal activity, it remains possible that the ostensibly ‘normal’ appearing LFP may possess some abnormal characteristics. We did not employ methods such as regression to mitigate differences between subjects and hippocampi, as we were concerned about potential biases of such regressions and due to the challenge of accurately quantifying the healthy stage of the hippocampi. To address temporal variations and abnormalities, we averaged over a high number of time-windows (500), with each group sampled from different recording times.

In estimating velocities, we employed Euclidean distances between electrodes as a proxy for transferred distances. However, it is important to note that the actual propagation distances are likely to be longer than the Euclidean distances for two primary reasons. Firstly, neural pathways do not follow straight lines, and secondly, while pathways are conventionally defined as unidirectional, they may only be effectively unidirectional with some uneven bidirectional flow. Consequently, the velocities we obtained may be underestimated.

## 4. Methods

### 4.1 Subjects

The study involved three subjects with drug-resistant epilepsy undergoing intracranial EEG monitoring, which included depth electrodes inserted along the hippocampal longitudinal axis as part of their pre-surgical evaluation. The study was approved by the Ethics Committees of the Hadassah Medical Center, Jerusalem, Israel.

The SEEG electrodes comprised 15 contacts, spaced 0.35 cm apart, with the most anterior contact positioned first (numbered 1). Each contact captured the local-field potential within a sphere around its tip, with a radius of approximately 5 mm. Table 1 provides a comprehensive overview of the subjects’ clinical, imaging, and electrophysiologic data.

**Table 1:** A comprehensive overview of the subjects’ clinical, imaging, and electrophysiologic data. Rt. Right, Lt. Left, MTS mesial temporal sclerosis, ASM ani seizure medication, PHT phenobarbital,LEV levetiracetam, CLB clobazam, CZP clonazepam, CBZ carbamazepine, LCM lacosamide, RNS responsive neurostimulation, ATL anterior temporal lobectomy, AH amygdalohypocampectomy.

|  | Pt. 1 Rt. hip | Pt. 1 Lt. hip | Pt. 2 Rt. hip | Pt. 2 Lt. hip | Pt. 3 Rt. hip |
| --- | --- | --- | --- | --- | --- |
| Atrophy | + (MTS) | - | - | - | + |
| Seizures | + | - | + | + | + |
| interictal activity | + | + | + | + | + |

**Table 1:** A comprehensive overview of the subjects' clinical, imaging, and electrophysiologic data.
|  | Pt.1 M | Pt.2 F | Pt.3 F |
| --- | --- | --- | --- |
| Age of onset/epilepsy duration (years) | 15/20 | 25/15 | 17/4 |
| Seizure semiology | Lt hand tingling, dialeptic, lt clonic | Anxiety, dialeptic | Lt. hand tonic, dialeptic |
| EEG | ii: Right and left temporal sharp waves<br>Ictal: diffuse onset, sometimes more pronounced right temporal | ii: Left temporal slowing and sharp waves<br>ictal: independent left and right frontotemporal onset | ii: right parietotemporal spikes<br>ictal: right posterior temporal |
| MRI | Right frontal cystic changes, right MTS | normal | Right frontal porencephaly, rt MTS |
| PET CT | Rt. frontotemporal hypometabolism | Rt. frontal and lt. mesial temporal hypometabolism | Right frontotemporal hypometabolism |
| Neuropsychological |  | Memory and executive functions impairments | Visual memory impairment, mild concentration difficulty and mild decrease in cognitive processing |
| FMRI language lateralization | Bilateral | Bilateral, left more than right | Left |
| Clinical hypothesis prior to SEEG | Independent bitemporal versus rt. temporal lobe epilepsy | Independent bitemporal seizures versus lt. temporal neocortical | Rt. hippocampal seizures, possible dual pathology |
| Clinical SEEG seizure onset | Rt. hippocampus. | Lt. temporal neocortex, rt. hippocampus. | Rt. hippocampus (24/27); unclear, probably rt. frontal (3/27 seizures) |
| ASMs | CBZ, CLB | CBZ, BRV, CLB, cannabis | CBZ, LVT, SOS clonazepam |
| Past ASMs | PHT, LEV | LTG, LEV, CZP, permapanel; VNS | LCM, VPA |
| Surgical intervention | Rt. ATL+AH | RNS (rt. and lt. hippocampus) | Rt ATL+AH |
| Pathology | Hippocampal sclerosis | N/A | Hippocampal sclerosis, focal cortical dysplasia IIIa |
| Post-procedure outcome | Seizure free | Seizure free | Seizure recurrence |

To ensure the accurate localization of electrode contacts within the hippocampus, a specialized neurosurgeon utilized merged CT/MRI images to identify and document all contact positions, distinguishing whether they were within or outside the hippocampal region. Supplementary figures 1-5 exhibit exemplar images derived from these merged CT/MRI images.

### 4.2 Software

For the bivariate (pair coherence) and the multivariate (four-node pathways) calculations, we used wavelet analysis with custom-developed software in IDL version 8.2.0 (Exelis Visual Information Solutions, Inc.). The wavelet software was provided by Torrence and Compo [30, 31]. Calculations were performed on 1sec long time-windows, each with 1024 points. The complex Morlet wavelet functions, which have been demonstrated to offer a favorable balance between time and frequency localization [18], were used. In the calculations, we set the smallest scale to 0.01, 1 millisecond for the time resolution and employed 31 scales to cover a frequency range of 7.2-96.8 Hz. This frequency range was selected to include the alpha to high gamma bands but excluded the theta frequencies due to the time windows’ length.

### 4.3 PLI Calculations

The Phase Lag Index (PLI) computation followed the methodology outlined by Stam et al [32, 33]. Initially, time-series (contact signals), each spanning 1 second with 1024 points, underwent wavelet transformation to transition into the time-frequency domain. Subsequently, pair phase coherence was calculated as a function of time and frequency for all possible 105 pairs. As spontaneous activity data is not synchronized to any specific event, the product of the wavelet coefficient of one time-series with the complex conjugate of the wavelet coefficients of the other time-series (that is the definition of coherence) was averaged over time. The pair coherence for the *i-j* pair was then calculated using the formula:

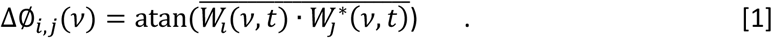

This process was iterated over 500 time-windows for each pair, and the phase coherence was determined for each window. PLI was subsequently calculated to define significant coherence for each pair over these 500 windows as:

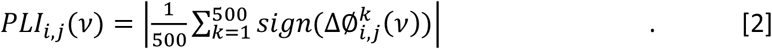

In addition, we computed the average phase-coherence and the average contact distance (in cm) over significant coherences (average over *l* significant pairs) for each frequency:

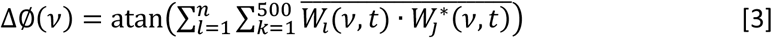

And

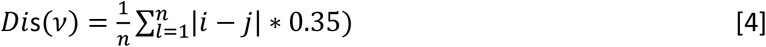

where *n* is the number of significant coherences at frequency *v*.

### 4.4 Pathway index (PWI) Calculations

In our prior study, we showcased the utilization of interactions among four coupled time-series to determine their temporal coupling order at each time-frequency point [12, 15-20]. Two key assumptions underpinned these inferences: firstly, that the coupling between the time-series was relatively weak, and secondly, that propagation was predominantly unidirectional. The fundamental analysis framework for the four coupled time-series has been previously outlined and will be briefly summarized here for clarity.

In our previous work, we demonstrated that the conditions for coherence between four time-series can be described by the mutual existence of three expressions, which we referred to as functional connectivity (FC) [6]:

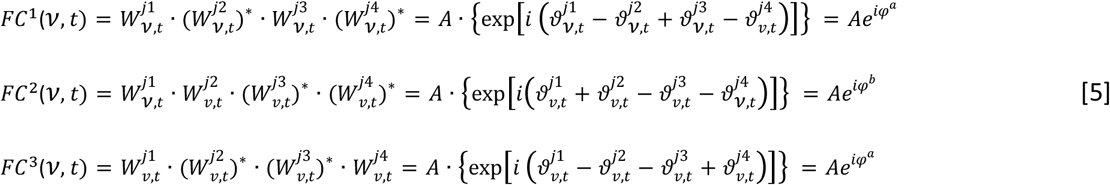

with, 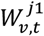and 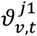 the wavelet coefficient and phase of time series ‘*j1*’ at a time-frequency point, where the numbers 1 to 4 represent the four time-series. A’ the amplitude, * denotes complex conjugate and ‘*i*’ the imaginary unit. Pair like phase-differences were expressed by the *φ*^*a*^, *φ*^*b*^, and *φ*^c^ phases as:

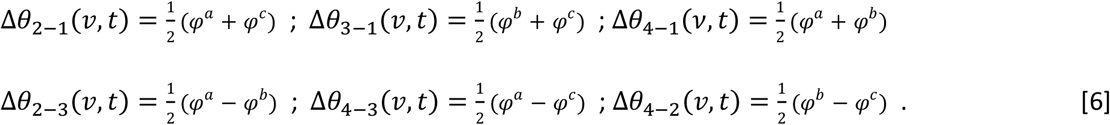

Similar to pair coherence for spontaneous activity data, we average the phases over time. These averaged phases are then utilized to determine the temporal order, referred to as pathways, at each frequency.

Our focus lies on effectively continuous and unidirectional pathways, indicating sequences that originate from one ensemble and sequentially pass through all other ensembles. For four-node pathways, there exist a total of 24 different pathways, as listed in Supplementary Table 1.

To establish pathways independent of the reference phase and the 2π limitation (as described in [9, 10], see example below), we selected a subgroup of pathways where all four phase differences, each with a different reference phase, yielded the same pathway that is, the same temporal order. Specifically, we defined a pathway’s index using the following criteria:

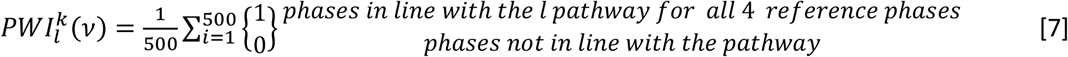

with ‘*l*’=1, 2… 24 corresponding to a pathway’s number in supplementary Table 1, *k* refers to the choice of the 4 time-series (out of the 1365 possibilities), and summation is over the 500 time-windows. We note that averaging of wavelet-coefficients over time-windows solves the intrinsic time-frequency uncertainty [30, 31].

For clarity, consider the following three pair-like phase-differences at frequency *v* : Δ*θ*_2_−1(*v*) = 50^°^, Δ*θ*3−1(*v*) = 8°^°^, Δ*θ*4−1(*v*) = 2°^°^. These coherences suggest the ‘1-4-2-3’ temporal order. In this case, the phase of the first time-series was used as a reference phase. The condition in Equation 7 ensures that the same time-order is obtained when all four phases are used as reference-phases, minimizing the risk of obtaining a phase difference that exceeds 2π. It is important to note that to obtain a directed pathway, we need to determine whether information flows from left to right or from right to left. For the purpose of the current study, where we exclusively utilize the average phase along the pathways (computed as 1⁄3 of the sum of individual phases (| Δ∅1−2| + | Δ∅2−3| + | Δ∅3−4|)), the directionality is not needed, only the temporal order calculated by equation 7.

Similar to the pair coherence calculations (Eq. 3 and 4), the average phase and distance along pathways are calculated over the 500 time-windows and for all significant pathways from the 1365 combinations, as a function of frequency.

### 4.5 Statistical analysis

We employed permutation non-parametric tests to compute the null distributions of PLI (Eq. 2) and PWI (Eq. 7) separately for each frequency. To ensure the uncoupling of the 15 contact signals, a random number generator was utilized to select time-windows from which the contact signals were obtained.

For the PLI calculations, data from subject 2 right hippocampus were used, and null calculations were repeated 10,000 times with PLI calculated on each iteration. For the PWI calculations, data from subject 1 left hippocampus were used, and this process was repeated 100,000 times with Equation 7 calculated on each iteration. Due to low null values in the PWI calculations, a higher number of iterations was needed.

In the null PLI calculations, a value of 0.19 corresponded to *p* < 1.6. 1°^−5^. To correct for multiple comparisons involving 105 * 31 combinations and frequencies, even using the most conservative Bonferroni correction, a PLI value of 0.19 corresponded to a corrected p-value of 0.05, and this value was used in the calculations.

For the PWI calculations, the highest value obtained for PWI was 0.057. The null distributions across all frequencies indicated that a PWI value of 0.05 corresponded to p<10-5. To account for multiple comparisons involving the 1,365 * 31 combinations of pathways and frequencies, we implemented a higher cutoff value. Since the null distributions could not estimate p-values for PWI values higher than 0.056 (even with 100,000 calculations), we set a conservative cutoff of 0.15. This cutoff value ensured that randomized null signals yielded zero pathways, while correct data resulted in only a few hundred pathways at specific frequencies.

## Acknowledgements

This work was funded by the Anges Ginges Foundation and the Prusiner-Abramsky 2021 award (GG).

## Data Availability

Data will be available upon request.

## Supplementary Material

In a separate file.

## Author Contributions

G.G.: Conception and design of the study, development of analyses, execution of analyses, and writing of the manuscript.

T.B.: Inspection of data, evaluating subject state and writing the manuscript.

Z.I.: Performing the surgery.

J.L: Evaluation of electrode and contact positions.

S.H: Evaluation of electrode and contact positions.

D.E.: Writing the manuscript.

## Notes

### Competing Interest Statement

The authors have declared no competing interest.

