## Supplementary material for "Velocities of Hippocampal Traveling Waves Proportional to Their Coherence Frequency": method

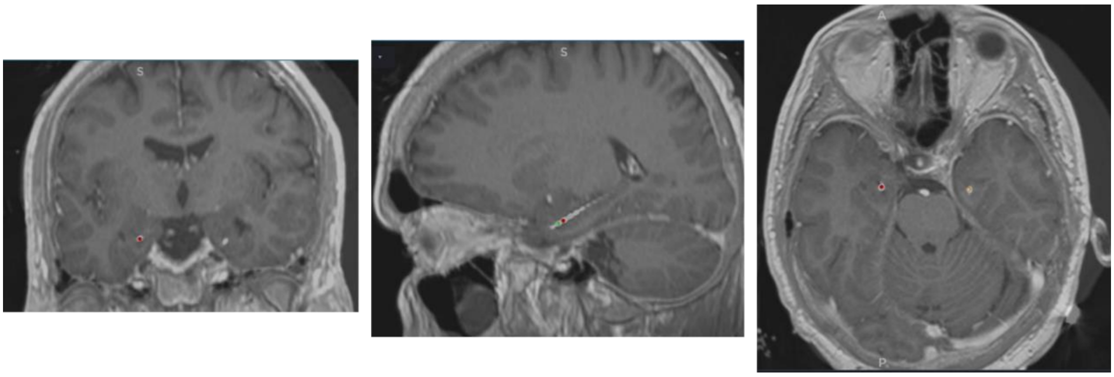


<u>Figure 1:</u> Merged MRI and CT Images of Subject 1, Left View. The red dot indicates contact number 5 on the SEEG electrode inserted to the left hippocampus.

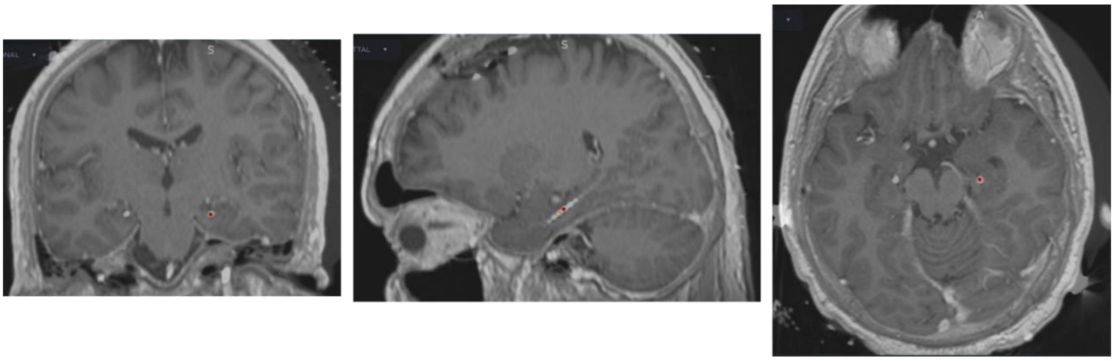


<u>Figure 2:</u> Merged MRI and CT Images of Subject 1, Right View. The red dot indicates contact number 5 on the SEEG electrode inserted to the right hippocampus.

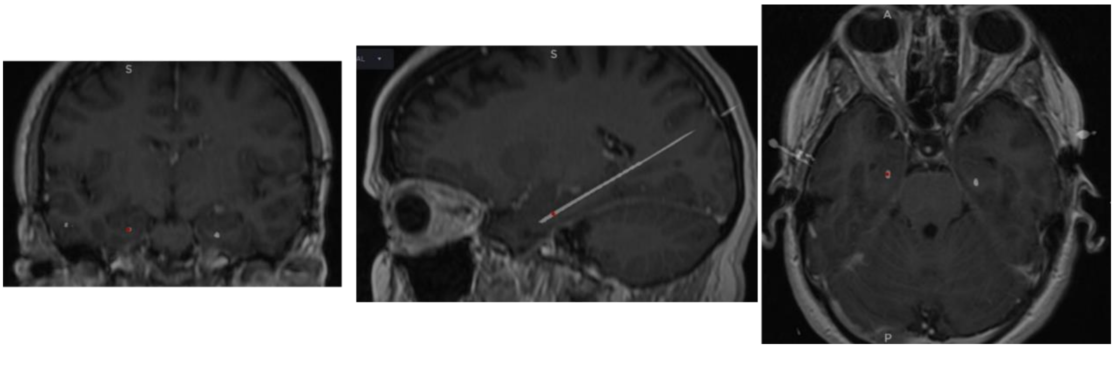


<u>Figure 3:</u> Merged MRI and CT Images of Subject 2, Left View. The red dot indicates contact number 4 on the SEEG electrode inserted to the left hippocampus.

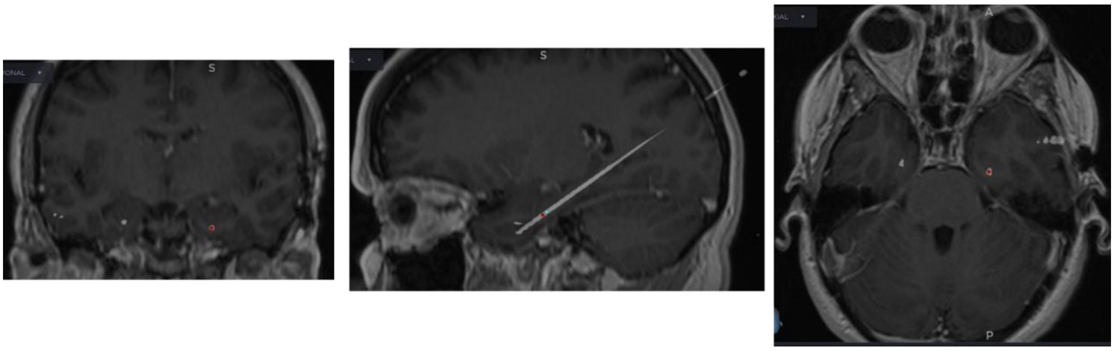

Figure 4: Merged MRI and CT Images of Subject 2, Right View. The red dot indicates contact number 5 on the SEEG electrode inserted to the right hippocampus.

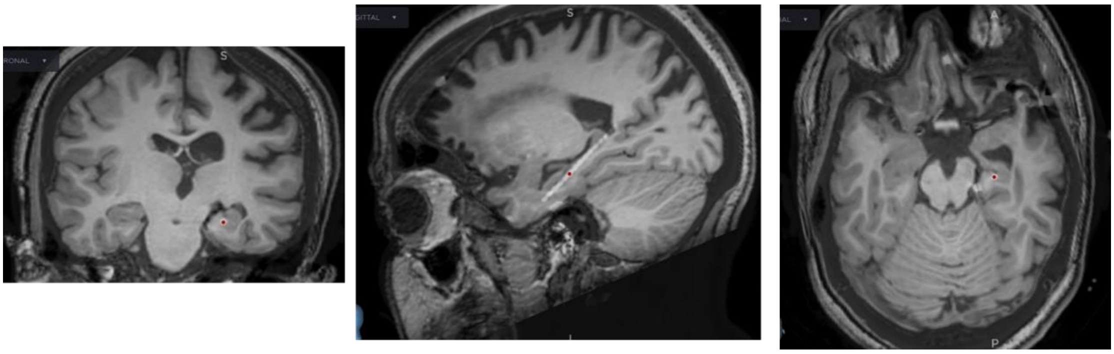

Figure 5: Merged MRI and CT Images of Subject 3, Right View. The red dot indicates contact number 6 on the SEEG electrode inserted to the right hippocampus.

**Table 1**

| PW number | Pathway |
|---|---|
| 1 | E1-E2-E3-E4 |
| 2 | E2-E3-E4-E1 |
| 3 | E3-E4-E1-E2 |
| 4 | E4-E1-E2-E3 |
| 5 | E1-E2-E4-E3 |
| 6 | E2-E4-E3-E1 |
| 7 | E3-E1-E2-E4 |
| 8 | E4-E3-E1-E2 |
| 9 | E1-E3-E4-E2 |
| 10 | E3-E4-E2-E1 |
| 11 | E4-E2-E1-E3 |
| 12 | E2-E1-E3-E4 |

| | |
|---|---|
| 13 | E1-E4-E3-E2 |
| 14 | E2-E1-E4-E3 |
| 15 | E3-E2-E1-E4 |
| 16 | E4-E3-E2-E1 |
| 17 | E1-E3-E2-E4 |
| 18 | E2-E4-E1-E3 |
| 19 | E3-E2-E4-E1 |
| 20 | E4-E1-E3-E2 |
| 21 | E1-E4-E2-E3 |
| 22 | E2-E3-E1-E4 |
| 23 | E3-E1-E4-E2 |
| 24 | E4-E2-E3-E1 |

**Table 1:** A list of all four-node pathways. E1 to E4 are the four contact signals.
